# SPACE: multimodal spatial CRISPR screening with whole-transcriptome readout at subcellular resolution in 3D models

**DOI:** 10.1101/2025.09.14.675819

**Authors:** Mengwei Hu, Yi Cui, Qianhui Huang, Khoi Chu, Sierra McKinzie, Michael Patrick, Sharanya Iyengar, Maerjianghan Abuduli, Marianne Spatz, Nandita Joshi, Brendan Miller, Shams Vellarikkal, Timothy Riordan, Danny Bitton, Jan Lubojacky, Iya Khalil, Federica Piccioni, Michael Rhodes, Alex Tamburino, Shanshan He, Joseph Beechem, Vanessa Peterson

**Author notes:** These authors contributed equally to this work (Mengwei Hu, Yi Cui).

## Abstract

Current spatial CRISPR screening technologies are limited by targeted readouts and high costs, restricting the scope of biological discovery. Here we present **SPA**tial **C**ell **E**xploration (SPACE), a spatial CRISPR screening platform that integrates whole-transcriptome profiling (∼18,000 genes), multiplexed protein detection (∼68 markers), and CRISPR perturbation mapping at subcellular resolution. SPACE significantly reduces whole-transcriptome profiling costs compared to sequencing methods while preserving spatial context. We demonstrate SPACE by screening 43 CRISPR knockouts (KOs) across ∼100,000 cells in hundreds of cancer-associated fibroblast (CAF)-tumor spheroids, obtaining whole-transcriptome and multiplexed protein readout from the same exact cells. SPACE revealed previously unknown regulatory mechanisms on tumor extracellular matrix (ECM) remodeling, and identified spatially-resolved ligand-receptor interactions and perturbation-specific spatial gene signatures that are not detectable with dissociation-based methods. This scalable, cost-effective platform provides a transformative framework for high-throughput spatial perturbation studies in complex tissue models.

## Introduction

Understanding how genetic perturbations affect cellular behavior within native tissue contexts is fundamental to drug discovery and precision medicine. While CRISPR screening has revolutionized functional genomics and emerged as a powerful tool for novel drug target identification, current approaches face limitations. Sequencing-based methods like Perturb-seq require tissue dissociation, destroying spatial information essential for understanding cell-cell interactions and tissue organization. Although recent technique advancements have led to multiple spatial CRISPR screen technologies that study different phenotypes^1–8^, they remain constrained by hypothesis-driven gene panels rather than enabling unbiased discovery afforded by whole-transcriptome profiling. Furthermore, most studies are restricted to 2D cultures, and no existing spatial CRISPR screen method simultaneously captures whole-transcriptome changes with spatial context at scale.

Here we present **SPA**tial **C**ell **E**xploration (SPACE), which overcomes these limitations by enabling whole-transcriptome spatial CRISPR screening at unprecedented scale and resolution. SPACE uniquely integrates three capabilities in a single assay: whole-transcriptome profiling (∼18,000 genes), CRISPR perturbation mapping, and high-plex targeted protein detection, while preserving subcellular spatial resolution in 3D models. This allows large-scale perturbation studies in complex model systems such as spheroids, assembloids, and organoids. Unlike dissociation-based methods that rely on computational inference, SPACE directly captures spatially resolved ligand-receptor interactions and cell-cell communication networks. This ability to simultaneously map perturbation effects, gene expression patterns, and cellular neighborhoods within intact tissues provides unprecedented insights into how genetic alterations reshape tissue architecture and intercellular signaling.

To demonstrate SPACE’s capabilities, we performed a comprehensive CRISPR screen across hundreds of CAF-tumor spheroids containing ∼100,000 cells, which would pose significant cost-related limitations for scaling with conventional sequencing approaches. SPACE achieved reliable technical performance, including confident CRISPR perturbation assignment, sensitive whole-transcriptome-scale RNA detection, and robust cell type classification. Our study allowed unbiased discovery of CAF-tumor biology from different aspects, including ECM remodeling, spatial ligand-receptor interaction and spatially variable genes. SPACE thus represents the first imaging-based platform to combine unbiased whole-transcriptome discovery with spatial CRISPR screening, offering a scalable and cost-effective framework for high-content cellular perturbation studies.

## Results

### SPACE successfully maps CRISPR perturbations in single cells in a cell pool

To establish SPACE for spatial CRISPR detection, we first validated the approach of detecting CRISPR guide RNAs (gRNAs) by *in situ* hybridization in 2D-cultured cells (Figure 1)^9, 10^. Because the gRNAs are ∼20-nt long, such short sequence only allows binding of 1 detection probe and may have suboptimal hybridization efficiency. To overcome this limitation, we paired each gRNA sequence with a 50-nt unique gRNA identifier (UGI) optimized for probe design criteria. We constructed lentiviral vectors expressing distinct gRNA-UGI pairs, transduced human SH-SY5Y-Cas9 cells individually to ensure each cell only receives 1 type of gRNA-UGI pair, then pooled the individually transduced cells for multiplexed CRISPR detection (Figure 1A). The lentivirus vector was modified from CROP-seq design to allow co-expression of gRNA and UGI in detectable mRNA transcripts (Figure 1B). The reliable expression of CRISPR tracrRNA and UGIs was confirmed via RNAscope (Figure 1C).

**Figure 1:**
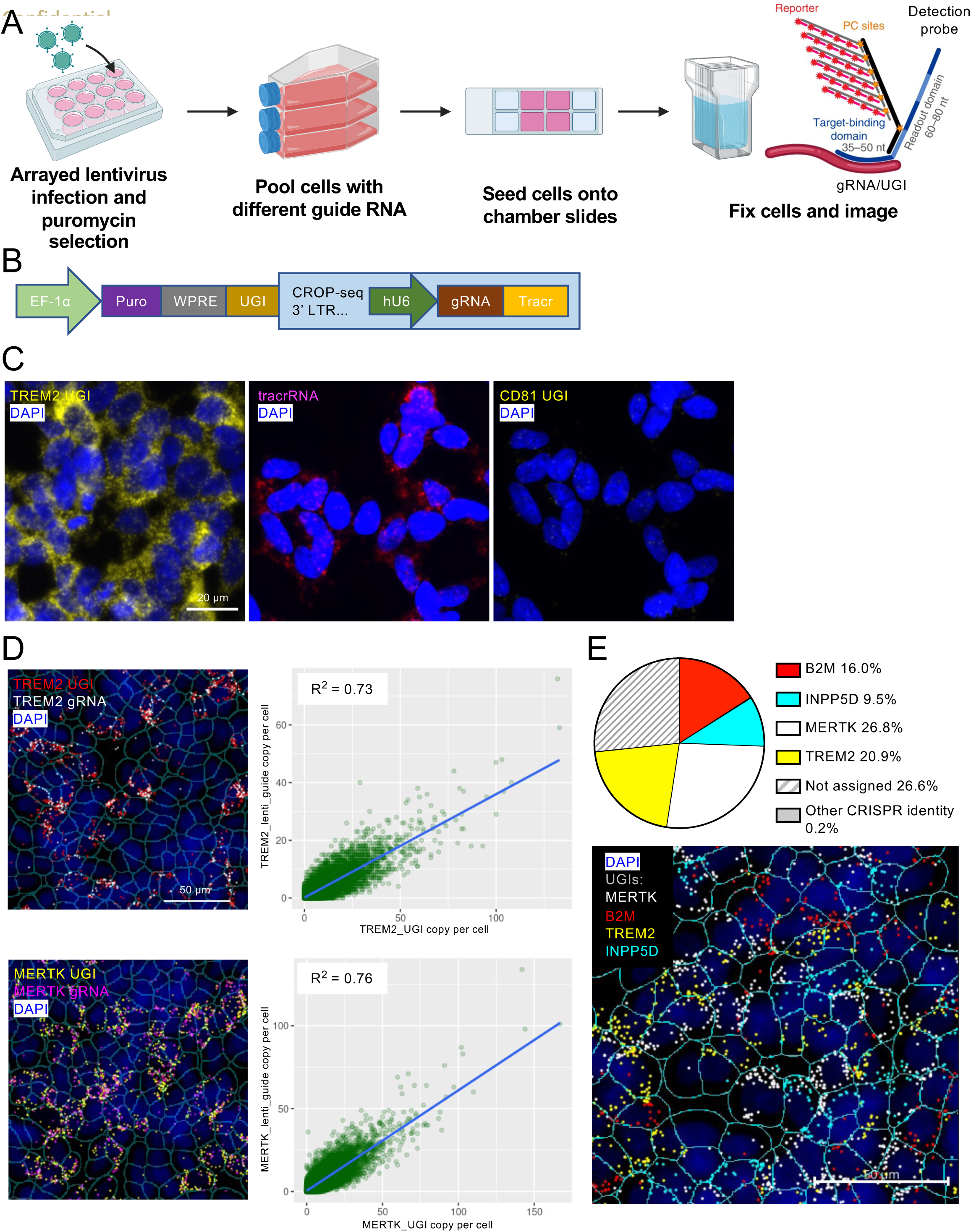
SPACE detection of CRISPR identities in a pool of lentivirus-infected cells. A: Diagram of experimental workflow. Cells are infected in separate wells by lentivirus that express different combinations of gRNAs and their corresponding UGIs, and selected for puromycin resistance. The cells expressing different gRNAs/UGIs are then pooled and seeded onto chamber slides, fixed, and continued with SPACE workflow. Diagram of SPACE probe and reporter for detection of CRISPR gRNAs and UGIs is shown (see also He *et al* Nat Biotech 2023, Khafizov *et al* Biorxiv 2024). Figure created with Biorender. B: Lentivirus vector design. The construct was modified from CROP-seq vectors. Briefly, the gRNA expression cassette (human U6 promoter – gRNA – tracr) is embedded in the 3’LTR, downstream of Puromycin resistance gene and WPRE that are driven by EF-1α promoter. A 50-basepair UGI is inserted between WPRE and 3’LTR. C: RNAscope detection of CRISPR molecules in cells expressing TREM2 gRNA and UGI. Left: detecting TREM2 UGI (yellow). Middle: detecting tracrRNA (magenta). Right: detecting CD81 UGI (yellow). Scale bar: 20 µm. D: SPACE detection of CRISPR molecules in mixed cell pools (left column) and correlation between the gRNA and UGI counts of the same gene (right column). Examples are shown for two gRNAs, TREM2 (up) and MERTK (down). Scale bar: 50 µm. E: Decoding CRISPR identity in mixed cell pool by SPACE-detected UGI. Up: Pie chart of cell decoding results. Down: SPACE image of UGI distribution in the cell pool. Scale bar: 50 µm.

The branched amplification strategy of the probe design scheme allows each target molecule to be tiled with multiple fluorescent reporters, generating high signal-to-noise ratios even for short transcripts like gRNAs and UGIs and allowing sensitive RNA detection (Figure 1A). In a cell pool of 4 different gRNA-UGI pairs, we detected on average 9.5 gRNA copies and 18.6 UGI copies per cell (n = 21,351 cells). The strong correlation between gRNA and UGI counts within individual cells (Figure 1D) demonstrated high detection specificity. Using a maximum expression assignment algorithm (see Methods), we confidently assigned CRISPR identity to 73.2% of cells (Figure 1E). These data establish that SPACE successfully decodes CRISPR identities to each single cell with high specificity in pooled cell populations.

### Establishing SPACE in 3D spheroid models

To enable SPACE technology in 3D models, we utilized a co-culture system of cancer-associated fibroblasts (CAFs) and tumor cells, a well-established model for studying CAF-tumor crosstalk and anticancer drug screening^11^. We CRISPR edited the CAFs targeting 42 different genes plus negative control *AAVS1* via arrayed electroporation, delivering gene-specific gRNA sets paired with unique UGIs to each well (Figure 2A). We co-cultured CRISPR-edited CAFs and unedited BxPC3 tumor cells in ultra-low attachment plates for 3 days to generate spheroids, which were subsequently fixed and arrayed in formalin-fixed paraffin-embedded (FFPE) blocks as tissue microarrays (microTMAs)^12^. This approach preserved the same plate map of CRISPR edit identity as the 96-well, providing ground truth for validation (Figure 2A). We further confirmed the spatial organization of CAFs and tumor cells via H&E and COMET staining using adjacent sections (Figure S1).

**Figure 2:**
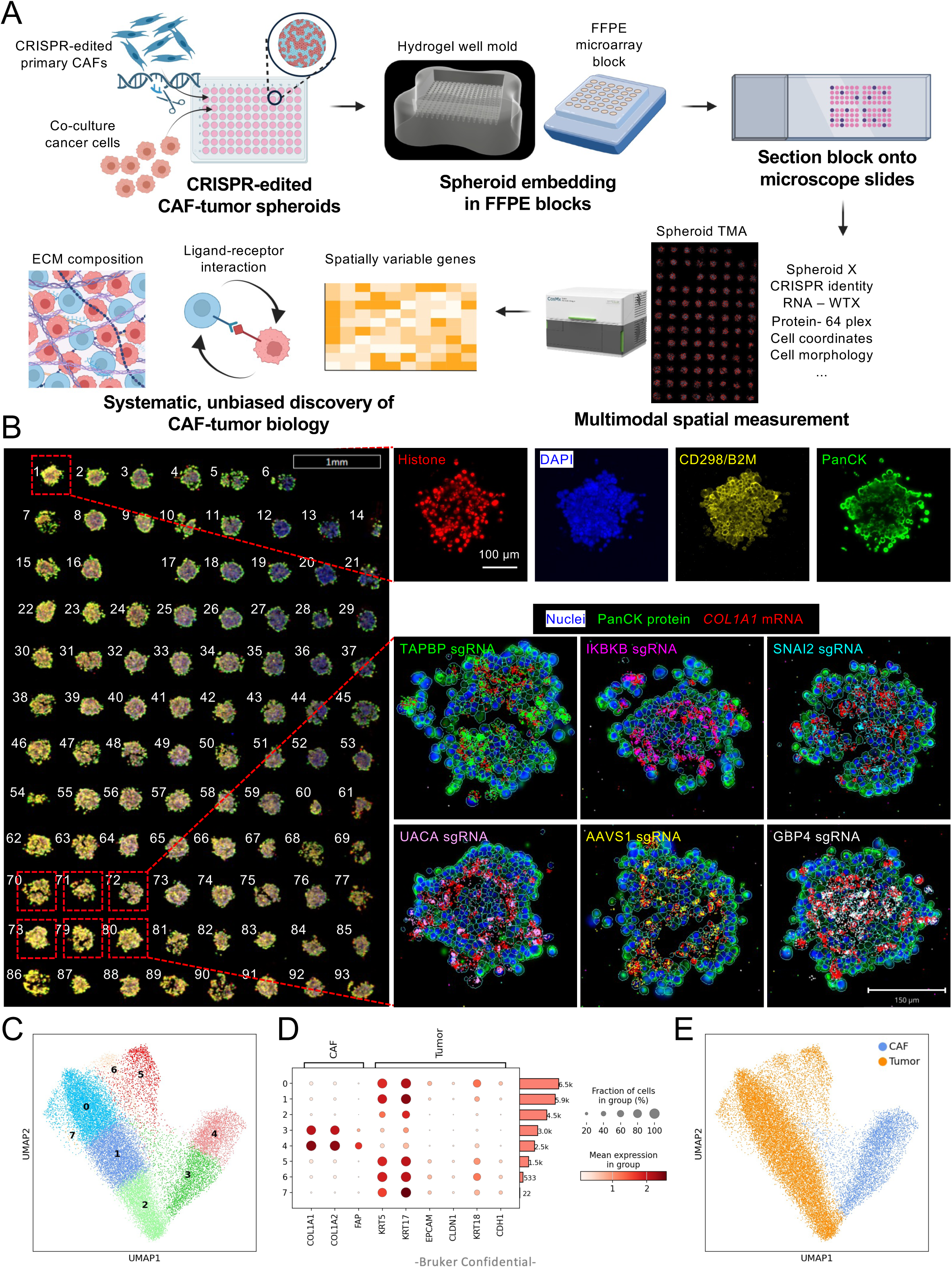
SPACE workflow in spheroid model. A: Diagram of SPACE experimental workflow. Primary cancer associated fibroblasts (CAF) are CRISPR edited, co-cultured with BxPC3 cancer cells in ultra-low attachment plates to allow spheroid formation, and fixed. The spheroids are then embedded in FFPE blocks while maintaining 96-well arrayed format for CRISPR identity confirmation. The sectioned slides undergo SPACE profiling of CRISPR molecules, transcriptome and protein on the same slide. Created in BioRender. B: Example images from SPACE workflow. Left: morphological staining of whole slide containing 93 spheroids placed in arrayed format. Scale bar: 1 mm. Top right: Morphological staining markers. Scale bar: 100 µm. Bottom right: composite images of representative spheroids, showing different synthetic gRNAs, *COL1A1* mRNA, PanCK protein and nuclear staining. Scale bar: 150 µm. C: Cell clustering using endogenous mRNAs detected by SPACE. D: Expression level of marker genes for CAF/tumor annotation in each cluster. E: Cell clustering with cell type of CAF and tumor annotated.

To distinguish different CRISPR perturbations, we designed detection probe panels targeting both gRNAs and UGIs, and coupled spatial CRISPR detection with either 6K-plex or whole-transcriptome (WTX) CosMx RNA profiling and protein markers (Figure 2B). A single slide containing ∼100 differentially perturbed spheroids yielded simultaneous detection of multiplexed endogenous transcripts, morphological markers, and CRISPR identifiers like gRNA/UGIs (Figure 2B). The comprehensive dataset offered by SPACE allows reliable annotation of CAF and tumor cell types (Figure 2C-E) and spatially resolved unbiased discovery on CAF-tumor interaction using multimodal perturbation data.

To evaluate SPACE assay performance in the spheroid model, we performed SPACE of 6K RNA + CRISPR gRNA/UGI on 2 slides with 93 spheroids per slide (Figure S2). The high CRISPR knockout efficiency in CAFs was confirmed using next-generation sequencing of CAF genomic DNA (Figure 3A). SPACE detection of endogenous mRNAs demonstrated exceptional reproducibility between two replicates when combining with CRISPR profiling (R² = 0.996, Figure S2A), detecting an average of 851 unique genes and 1,445 transcripts per cell (n = 45,255 cells). For CRISPR detection accuracy, we quantified gRNA and UGI counts in CAFs and ranked them by abundance. The top-ranked gRNA and UGI in each spheroid consistently matched the expected CRISPR, far exceeding background signal from off-target detection (Figure S2B and C). This high signal-to-noise ratio allowed confident perturbation assignment using a maximum expression algorithm for each spheroid.

**Figure 3:**
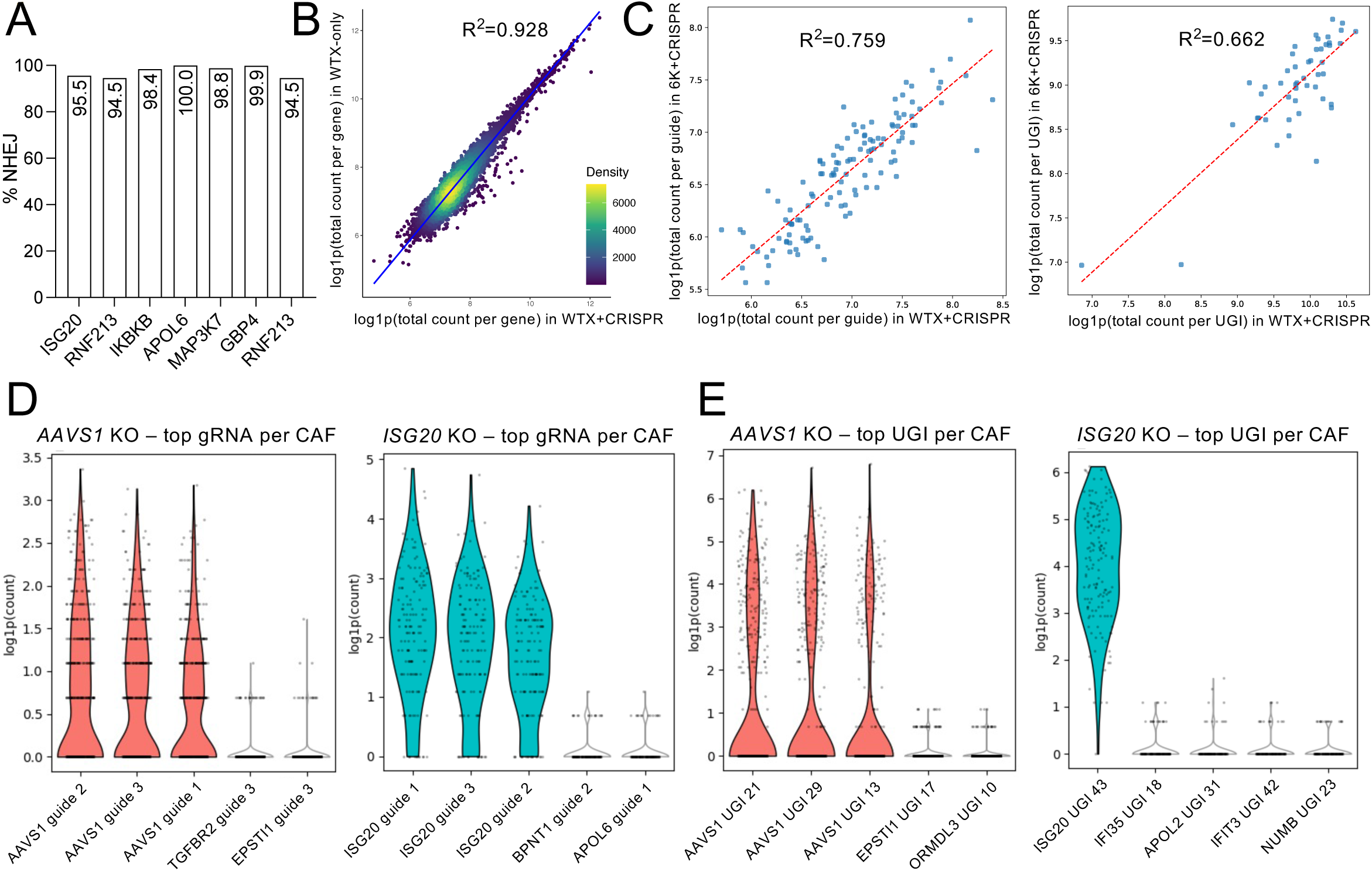
SPACE assay performance with WTX RNA panel. A: CRISPR editing efficiency in select CAF KO cells. B: Gene expression correlation between WTX+CRISPR experiment (x axis) and WTX-only experiment (y axis). C: Correlation of detected CRISPR molecule count between WTX+CRISPR (x axis) and 6K+CRISPR (y axis). Left: gRNA count correlation; right: UGI count correlation. D: Counts per CAF cell of the top 5 most abundant gRNAs in selected spheroid. Left: *AAVS1*-KO spheroids; right: *ISG20*-KO spheroids. E: Counts per CAF cell of the top 5 most abundant UGIs in selected spheroid. Left: *AAVS1*-KO spheroids; right: *ISG20*-KO spheroids.

### SPACE whole-transcriptome assay compatibility

We further asked if SPACE could scale to WTX profiling while maintaining CRISPR detection fidelity. By performing SPACE using CosMx WTX RNA panel with or without CRISPR molecule profiling, we found minimal impact on transcriptome coverage when incorporating CRISPR detection in SPACE (Figure 3). The WTX+CRISPR experiment detected an average of 1,518 unique genes and 2,188 transcripts per cell (n = 24,548 cells), which is highly comparable to an average of 1,516 genes and 2,405 transcripts per cell with WTX-only (n = 21,192 cells), and the transcript counts per cell is highly correlated between the two experiments (R^2^=0.928, Figure 3B). Compared with 6K+CRISPR experiment, WTX+CRISPR increased gene detection by 78% and transcript detection by 51%, demonstrating successful scaling of SPACE to WTX coverage.

CRISPR detection remained highly robust across different RNA panel complexities, with strong correlations between 6K+CRISPR and WTX+CRISPR experiments for both gRNA (R^2^=0.759) and UGI counts (R^2^=0.662) (Figure 3C). In WTX+CRISPR, the most abundant gRNA or UGI in CAFs also matched the ground truth with high signal over background (Figure 3D and E), ensuring confident CRISPR identity assignment. With refined cell segmentation (Figure S3), we confidently identified CAF and tumor populations based on the expression levels of marker genes, such as collagen genes and *FAP* for CAF lineage^13^, and *KRT5* and *KRT17*^14^ for tumor lineage (Figure 2C-E and Figure S3D). Collectively, these results establish SPACE as a robust platform for simultaneous CRISPR perturbation mapping and transcriptome profiling in 3D models, with scalability from targeted panels to whole-transcriptome coverage without compromising detection accuracy.

### SPACE reveals ISG20 as a novel regulator of MMP pathway activity

The high-dimensional spatial perturbation data generated by SPACE enabled unbiased discovery of gene regulatory networks in CAF-tumor interplay. By analyzing differentially expressed genes between different CAF gene KOs compared to *AAVS1*-KO, we found that *ISG20*-KO in CAFs significantly reduced the expression level of multiple matrix metalloproteinase (MMP) family genes in CAFs while simultaneously upregulated MMP inhibitor genes (Figure 4A-C). Reactome Pathway analysis confirmed systematic depletion of MMP activation processes in *ISG20*-KO spheroids (Figure 4D). We further validated MMP1 downregulation in *ISG20*-KO CAFs via both RNAscope (Figure 4E) and immunofluorescence (Figure 4F). These data suggest that SPACE robustly detects transcriptome changes caused by CRISPR edits, enabling confident discovery of novel biology at scale. Given that MMPs are crucial in remodeling ECM and regulating tumor progression and metastasis^15^, this finding has significant therapeutic implications. Notably, high ISG20 expression in tumor is associated with worse patient survival (Figure S4) in multiple cancer types, underscoring the clinical relevance of this finding. While MMPs are implicated in various cancer and thus is a popular therapeutic target, to date there is only one FDA-approved MMP inhibitor^16^. Our findings suggest that ISG20 inhibition may be an alternative therapeutic strategy to modulate MMP pathway activity, and mechanistic investigation of the ISG20-MMP regulatory axis is needed. Taken together, our results demonstrate the great potential of SPACE in translational research and drug discovery.

**Figure 4:**
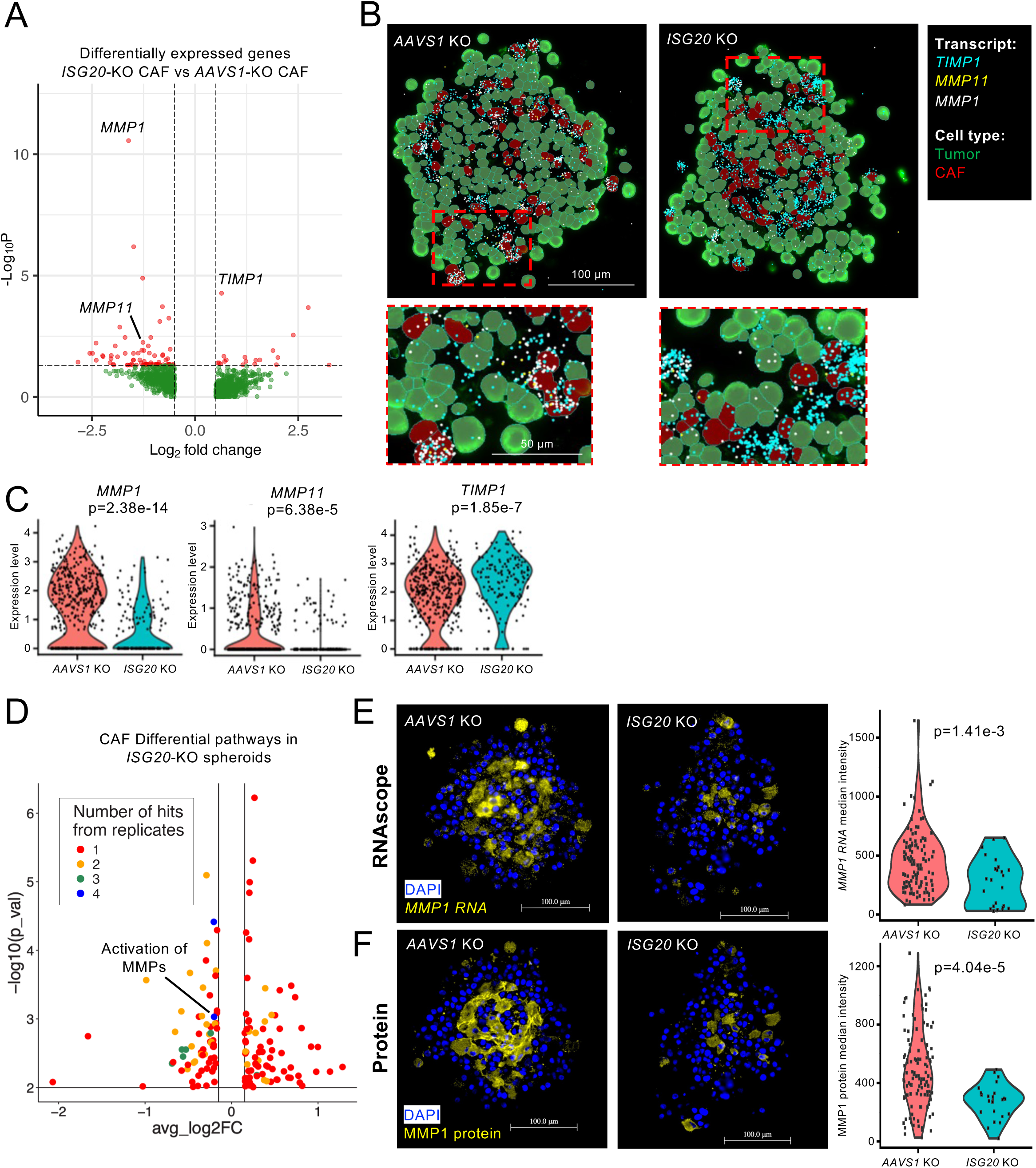
I*S*G20 KO led to downregulation of MMP pathway in CAFs. A: Volcano plot of differentially expressed genes of CAFs in *ISG20*-KO vs *AAVS1*-KO spheroids with selected MMP pathway-related genes labeled. B: Representative images from WTX+CRISPR SPACE showing selected RNA species in *AAVS1*-KO (left) and *ISG20*-KO (right) spheroids. Cell types are color-coded. Scale bar: 100 µm; 50 µm (zoom-in). C: Violin plots of selected gene expression level in CAFs in *AAVS1*-KO and *ISG20*-KO spheroids quantified from WTX+CRISPR SPACE. P values were calculated by Wilcoxon rank sum test. D: Reactome pathway analysis of *ISG20*-KO spheroids compared to *AAVS1*-KO spheroids. E: RNAscope detection of *MMP1* transcripts in *AAVS1*-KO and *ISG20*-KO spheroids using adjacent sections. Representative images and violin plot of CAF (represented by PCK-/Vimentin+) *MMP1* RNA median intensity are shown. P value was calculated by Wilcoxon rank-sum test. Scale bar: 100 µm. F: Multiplexed immunofluorescence detection of MMP1 protein in *AAVS1*-KO and *ISG20*-KO spheroids using adjacent sections. Representative images and violin plot of CAF MMP1 protein median intensity are shown. P value was calculated by Wilcoxon rank-sum test. Scale bar: 100 µm.

Tumor cell density has been shown to regulate various cancer behaviors such as metastasis^17^. The effect of cell density on gene expression caused by gene perturbations has also been observed in other spatial CRISPR screens, such as Perturb-FISH^4^. To investigate whether local tumor cell density modulates the MMP pathway activity, we categorized CAF cells into low-tumor-density and high-tumor-density groups based on their neighboring tumor cell counts (see Methods), and compared their MMP pathway activity across different perturbations (Figure S5). In the control *AAVS1*-KO spheroids, low-tumor-density CAFs exhibited significantly higher MMP pathway activity than high-tumor-density CAFs (Figure S5A and B). However, *ISG20*-KO in CAFs abolished this density-dependent gradient (Figure S5C and D). This spatial heterogeneity in perturbation response demonstrates SPACE’s capability to reveal how genetic perturbations reshape tissue microenvironment interactions. Our findings also highlight the critical importance of preserving spatial context when studying gene function in complex tissues.

### Spatial mapping of ECM-mediated CAF-tumor interactions

The ECM is an important and integral component of tumor microenvironment and mediates the crosstalk between CAF and tumor^13^. ECM proteins, such as fibronectin and collagen, can bind to receptors on tumor cell surface and regulate cell growth and migration^18^. We suspected that the CRISPR edits in CAFs may reshape ligand-receptor (LR) interactions at the CAF-tumor interface and thus regulate tumor cell behavior. Leveraging the spatial resolution offered by SPACE, we refined LR interaction analysis using physically adjacent cells, as they have higher likelihood of biologically meaningful physical LR interactions and thus will increase our confidence in finding true LR interactions. For each perturbation, we calculated the LR interaction with or without spatial information, treated non-spatially-informed LR as the background interaction level, and subsequently derived the enrichment/depletion of LR interactions at CAF-tumor boundaries (Figure 5A). Notably, *ISG20*-KO spheroids displayed increased interaction between CAF collagens and tumor CD44 (Figure 5A and B), potentially related to its reduced MMP pathway activity and consequently less collagen degradation. We also found that *RNF213*-KO spheroids display stronger interaction between CAF and tumors through various ECM proteins and cell adhesion molecules, including collagen, fibronectin, integrins and SDC4 (Figure 5A and B). Consistent with prior reports that strong ECM engagement through integrins can activate proliferative signaling pathways in tumors^18^, tumor cells in *RNF213*-KO spheroids exhibited upregulated GO terms associated with cell division and proliferation, including chromosome segregation and nucleosome assembly and organization (Figure 5C).

**Figure 5:**
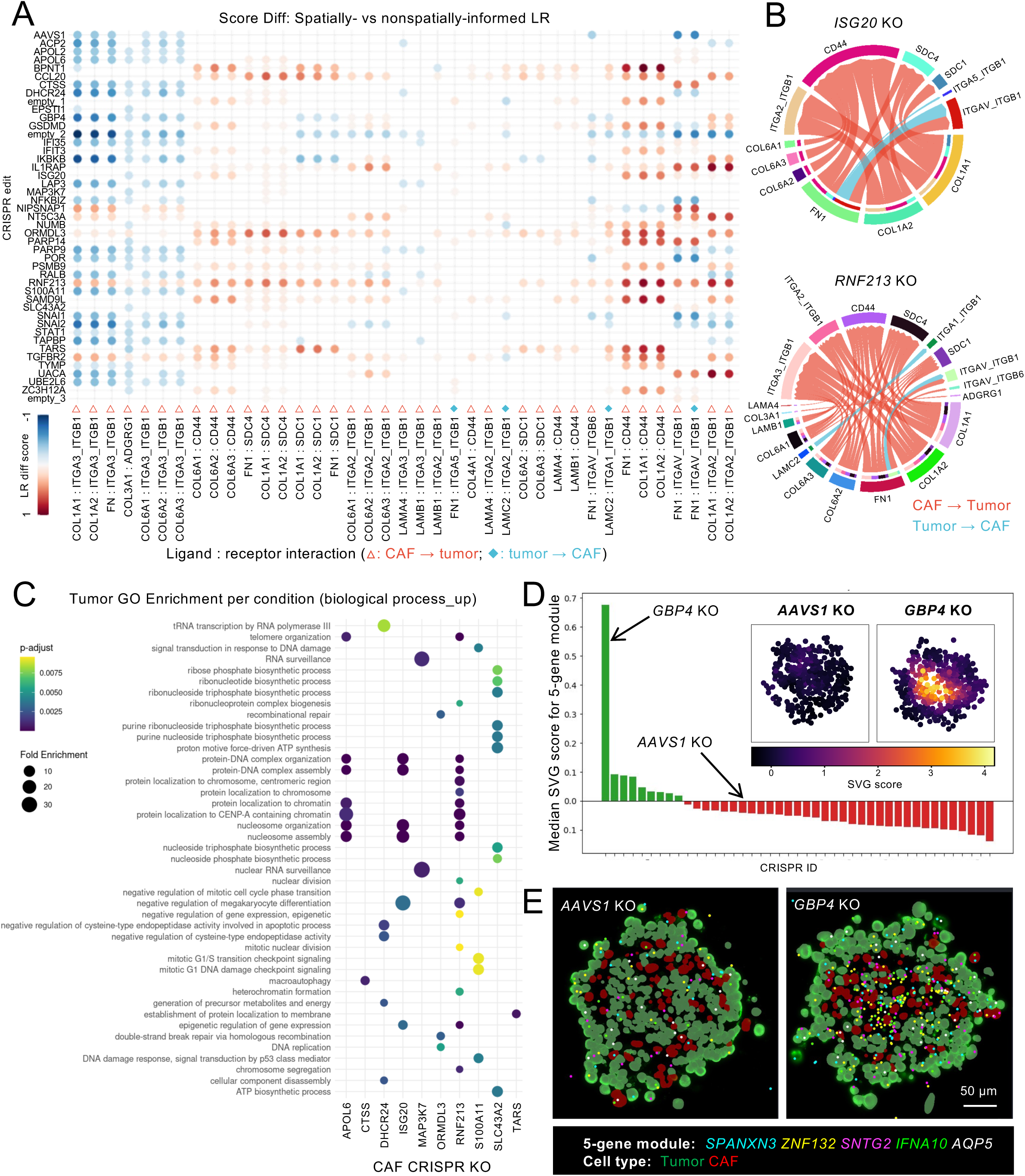
Spatially resolved ligand-receptor interaction and spatially variable gene module in CAF spheroids with different gene KO. A: LR score differences between the spatially informed vs nonspatially informed analysis for each CRISPR edits (blue – high LR interaction score in nonspatially informed analysis; red – high LR interaction score in spatially informed analysis). Only LR pairs identified in more than 5 KO conditions are presented. In x axis labels, the LR pairs are separated by “:”, with ligands before the colon and receptors after the colon. Red outline triangles indicate CAF-to-tumor interactions, and blue solid diamonds indicate tumor-to-CAF interactions. B: Chord diagrams showing the spatially informed LR pairs specific to *ISG20*-KO and *RNF213*-KO spheroids compared to nonspatially informed analysis (LR pairs with score difference > 0). The arc length of each LR pair represents its weight among all identified LR pairs. C: Enriched GO terms (biological process) of upregulated tumoral genes in selected CRISPR-edited spheroids. D: Waterfall plot of median SVG score for the 5-gene signature in each CRISPR edits and SVG score heatmap of selected spheroids. E: Representative images from SPACE showing the RNAs of the 5-gene SVG in *AAVS1*-KO and *GBP4*-KO spheroids. Scale bar: 50 µm.

SPACE enabled unbiased identification of spatially variable genes (SVGs) that are specific to certain gene edits, which indicate novel gene regulatory network in mixed cell population and are invisible to sequencing-based approaches. By performing SVG analysis, we found a spatially coordinated 5-gene signature in *GBP4*-KO spheroids (Figure 5D and E). *GBP4* is an interferon-stimulated gene that can negatively regulate type-I interferon signaling^19^, a pathway with complex roles in cancer^20^. Interestingly, GBP4 is among the most abundant GBP family proteins in pancreatic cancer, and is associated with poor prognosis in pancreatic cancer patients^21^. Within this gene signature, *IFNA10* is a member of type-I interferon, while *AQP5* and *ZNF132* have been reported as potential prognostic markers in multiple cancer types^22, 23^, suggesting a potential link between GBP4 and an interferon-associated spatial signaling program. While further investigation is needed to dissect the relationship between *GBP4* and the 5-gene signature, these findings demonstrate SPACE’s ability to reveal how single-gene perturbations propagate through tissue space to reshape multicellular communication networks.

### Multimodal SPACE uncovers EMT regulation in tumor microenvironment

Lastly, to further unleash the multimodal potential of SPACE and thoroughly utilize the rich phenotypic landscape offered by spatial CRISPR screening, we developed an integrated workflow combining whole-transcriptome profiling (∼18,000 genes), CRISPR perturbation mapping, and highly multiplexed protein detection (68 markers) on a single tissue section—the highest phenotypic dimensionality achieved in spatial CRISPR screening to date (Figure 6 and Table S1). We first captured protein signal using oligo-conjugated antibody and sequential FISH, and then removed the antibody staining and proceeded with the same SPACE RNA profiling workflow (see Methods) (Figure 6A). This multimodal approach simultaneously resolved distinct spatial patterns across molecular modalities within individual spheroids: endogenous transcripts, CRISPR identifiers (gRNA/UGI), and protein markers each revealed complementary aspects of CAF-tumor organization (Figures 6B-D). Similar to the WTX+CRISPR experiments, the multimodal SPACE successfully identified CAF and tumor populations (Figure 6E and F). By integrating genome-wide transcriptional profiling with targeted proteomics and perturbation mapping, multimodal SPACE enables hypothesis-free discovery while comprehensively capturing perturbation effects across multiple biological scales.

**Figure 6:**
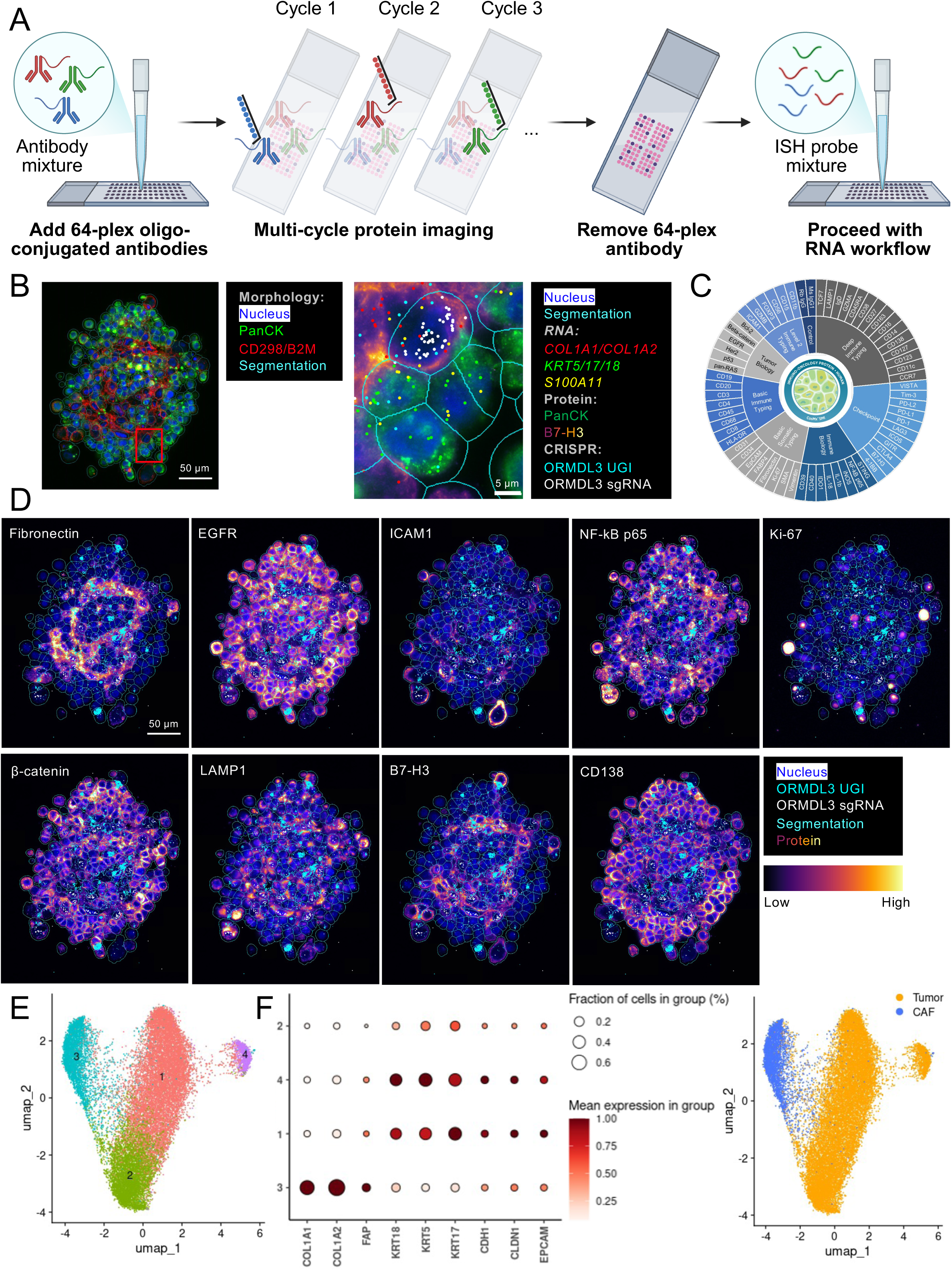
Multimodal SPACE. A: Diagram of multimodal SPACE workflow. Samples were first stained with oligo-conjugated antibody mixture and imaged on CosMx for 64-plex protein detection. The protein staining were then removed (see Methods) and the samples were further processed with SPACE WTX workflow for RNA profiling. Created in Biorender. B: Representative images of RNA detection of multimodal SPACE. Selective transcripts and morphological markers for cell segmentation are shown for the same *ORMDL3*-KO spheroid. Scale bar: 50 µm; 5 µm (zoom-in). C: 64-plex protein panel and their key biological purposes. D: Representative images of protein detection of multimodal SPACE. Selective protein markers are shown for the same spheroid as B. Scale bar: 50 µm. E: Cell clustering of multimodal SPACE based on endogenous mRNAs. F: Expression level of selected marker genes across different cell clusters and cell type annotation of multimodal SPACE.

Epithelial-to-mesenchymal transition (EMT) has been heavily implicated in multiple stages of tumor progression, including metastasis, drug resistance mechanisms, and CAFs have been found to be a key inducer of tumor cell EMT^24^. With multimodal SPACE data, we investigated how different CAF gene edits influence the EMT process in tumor cells. We computed single-cell EMT scores for tumor cells by integrating protein staining intensity of canonical EMT markers: vimentin, SMA, and fibronectin (mesenchymal) versus EpCAM (epithelial) (see Methods). We observed a significant negative correlation between the tumor EMT score and their distance to nearest CAFs (Figure S6), confirming CAFs as spatial EMT inducers. We then ranked the tumor EMT score for each CAF gene edit from low to high (Figure S7A), and visualized tumor cell EMT scores in a spatially resolved manner (Figure S7B). To validate that protein-derived EMT scores are in accordance with transcriptional reprogramming, we performed differential gene expression analysis between EMT-high and EMT-low tumor cells (Figure S7C). As expected, *VIM*, a canonical EMT marker gene^25^, was strongly upregulated in EMT-high cells. Prior studies have shown that tumor cell EMT is characterized by downregulation of epithelial lineage genes (including keratins) and increased ECM degradation via upregulation of MMP family members^25^.

Consistent with these reports, in EMT-high tumor cells, we observed downregulation of keratin genes such as *KRT13* and *KRT19* and upregulation of MMPs and collagens genes. This demonstrates that multimodal SPACE can capture molecular changes across multiple data modalities from the same cell, providing orthogonal layers of information that enhance our understanding of underlying biological mechanisms. By integrating protein and RNA measurements, this approach offers a generalizable framework for multiomic spatial CRISPR screens that can be applied across diverse models and disease contexts.

## Discussion

Here, we introduce SPACE, a scalable and versatile spatial CRISPR technology. We performed rigorous quality assessments of SPACE data, validated the biological insights it enabled using orthogonal approaches, and leveraged SPACE to perform hypothesis-free, data-driven discovery in a clinically relevant model. SPACE is the first spatial CRISPR screening platform to integrate whole-transcriptome profiling, highly multiplexed protein detection and CRISPR identity mapping at subcellular resolution in 3D models, achieving the highest phenotypic dimensionality in spatial CRISPR screening to date. By analyzing hundreds of thousands of cells across hundreds of CAF-tumor spheroids, we demonstrate several advantages of SPACE over existing technologies (Table S1) including compatibility with FFPE-embedded complex 3D models, preservation of spatial neighborhood information, and the ability to enable hypothesis-free discovery across a broad phenotypic landscape. Together, these features establish SPACE as a transformative advance in functional genomics and high-throughput screening.

### Technical advantages and scalability

The SPACE workflow confers several unique advantages. First, our imaging-based screen with whole-transcriptome readout profiled close to 100 spheroids per slide, with room to scale depending on spheroid or organoid size. For spheroids or organoids up to ∼500 µm, it is possible to fit >1,000 per slide, providing substantial potential for high-throughput screening. This increase in screening throughput also reduces the cost per cell/ per spheroid/organoid, making large-scale perturbation studies in 3D models economically feasible. Second, SPACE’s compatibility with FFPE samples expands its potential applications to patient-derived organoids and clinical specimens. This opens new avenues for precision medicine and personalized functional genomics, enabling patient-specific tissues to be screened for optimal therapeutic strategies. Third, the platform’s multimodal capabilities capture phenotypic changes at high resolution, allowing a single biological question to be addressed from multiple perspectives and providing orthogonal validation within the same experiment. For example, in our multimodal EMT analysis, protein-based EMT scores were in accordance with transcriptional signatures, increasing confidence in findings and highlighting how EMT is regulated at multiple molecular layers.

### Novel, spatially-resolved biological insights

SPACE revealed previously unknown regulatory mechanisms and exemplified the importance of preserving tissue architecture in perturbation studies. Using SPACE, we uncovered a novel regulatory role for ISG20 in MMP signaling within CAFs, suggesting ISG20 as a potential therapeutic target for modulating the ECM. Spatially-resolved ligand receptor (LR) analysis enabled us to dissect biologically meaningful LR interaction in a spatially informed manner, increasing confidence in the identified LR relationships. Our identification of RNF213 as a regulator of ECM-mediated cancer proliferation demonstrates how spatial screening can reveal functional connections between genotypes and phenotypes, providing mechanistic insight into how CAFs shape tumor behavior through ECM remodeling. Lastly, we discovered novel spatially variable gene (SVG) patterns specific to certain gene knockouts in CAFs, potentially representing previously unrecognized gene regulatory networks. Collectively, these findings illustrate how incorporating spatial information into functional genomics facilitates the study of gene functions in disease-relevant contexts, helping to elucidate the mechanisms of action for therapeutic targets and deepening our understanding of disease biology.

### Broad implications for drug discovery and precision medicine

SPACE addresses a critical gap in the drug discovery process by enabling functional and mechanistic interrogation of genes within physiologically relevant tissue contexts. Traditional drug screening in 2D cultures often oversimplifies the complex disease biology and fails to capture the cell-cell interactions that govern therapeutic response *in vivo*. SPACE addresses this gap by enabling systematic perturbation analysis in 3D models that more faithfully recapitulate disease context.

Although we showcase SPACE application in cancer biology, the same methodology can be broadly applicable to different disease area and organoid models. For example, SPACE could be applied to brain organoids to study gene functions in cell-cell communication, providing insights into psychiatric and neurodegenerative diseases. In immunology, SPACE may enable investigation of gene functions underlying immune cell infiltration, activation, and therapeutic response in autoimmune diseases and cancer immunotherapy. More broadly, the comprehensive perturbation datasets generated by SPACE across diverse tissue contexts will be a valuable resource for generative AI and machine learning models to learn, predict and unify causal principles of human biology^3^.

### Limitations and future directions

Several aspects of SPACE can be further improved to broaden its applicability. While our screen demonstrates feasibility, the number of gene perturbations accommodated by SPACE can be expanded. Achieving this will require optimization on gRNA or UGI capture probe design, reductions in on-instrument imaging time, and continued development of computational infrastructure to support large-scale data storage and analysis. The current SPACE study utilizes constitutive CRISPR knockouts, which limits its utility for studying essential genes or genes with functional redundancy. This limitation could be addressed by expanding SPACE to inducible CRISPR systems or CRISPRi and CRISPRa approaches.

Beyond genetic perturbations, UGI barcoding could also be paired with chemical perturbations, enabling high-throughput, spatially resolved compound profiling in organoid models. Such extensions would facilitate systematic studies of drug mechanism of action and further accelerate drug discovery efforts.

### Conclusions

In summary, SPACE represents a paradigm shift in functional genomics, moving beyond dissociated single-cell analysis in 2D toward spatially-resolved investigation of gene function in 3D, with unprecedented breadth and depth of phenotypic characterization. As the field increasingly embraces complex models that better recapitulate human physiology and disease, we anticipate that SPACE will accelerate causal, mechanistic insight into gene function in the disease-relevant context, ultimately enabling facilitating the development of more effective therapeutic strategies.

**Figure S1:**
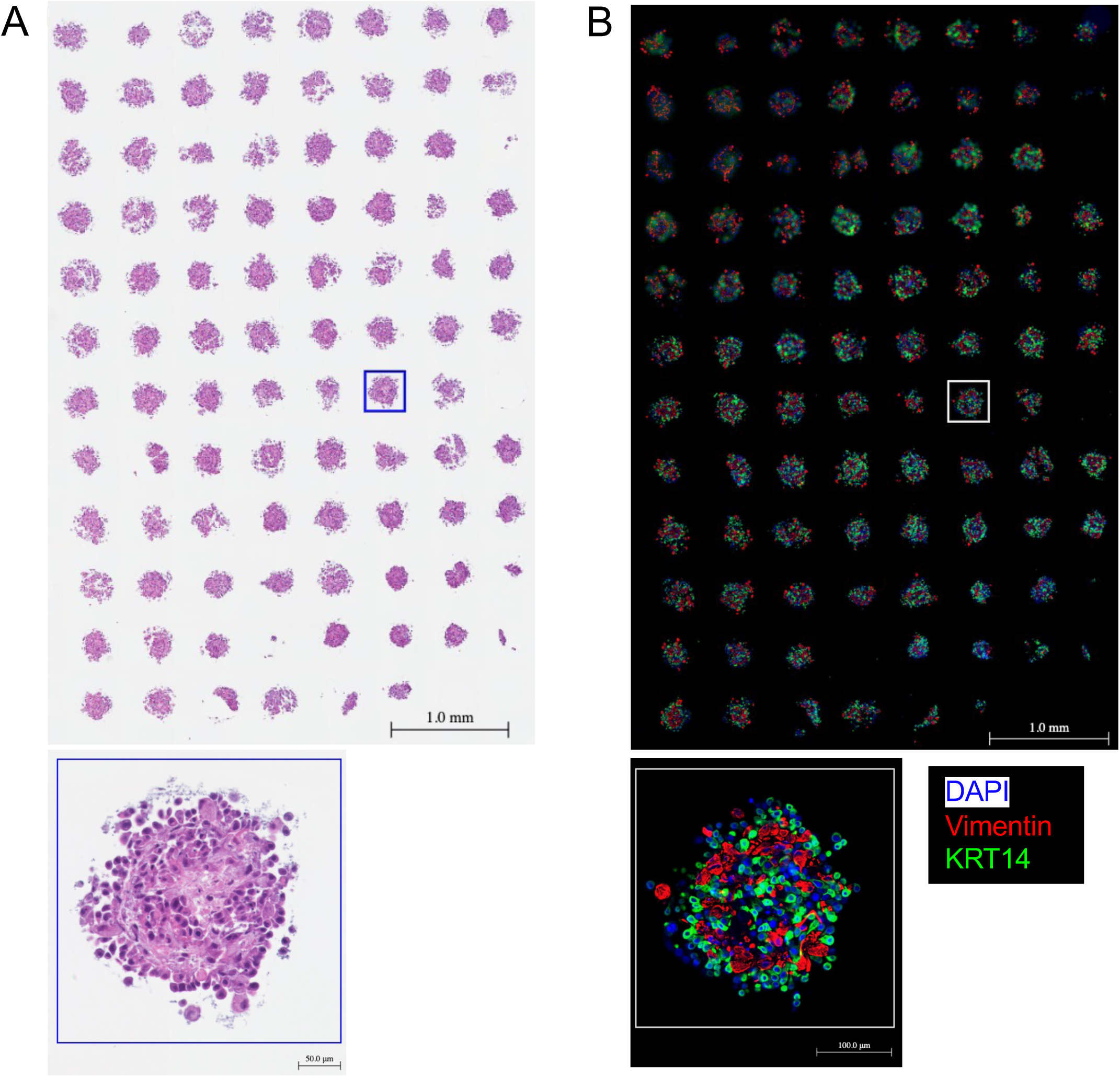
H&E and COMET staining of SPACE spheroid array. A: H&E staining. B: COMET protein staining showing selected markers: DAPI, Vimentin and KRT14.

**Figure S2:**
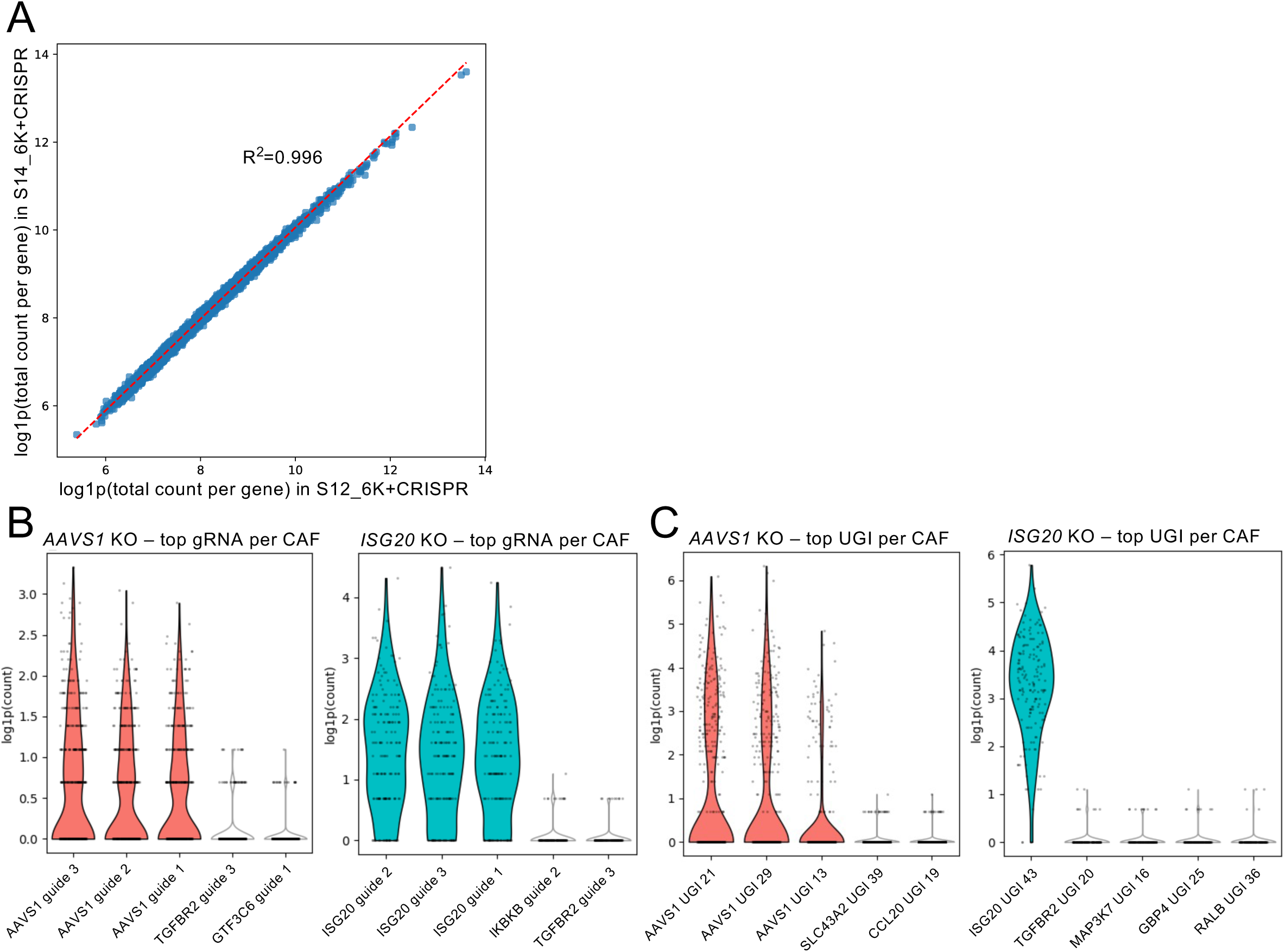
SPACE assay performance with 6K RNA panel. A: Gene expression correlation between two slides in a 6K+CRISPR SPACE assay. Each dot represents the log-transformed (log1p) total count of the gene within the dataset. B: Counts per CAF cell of the top 5 most abundant gRNAs in selected spheroid. Left: *AAVS1*-KO spheroids; right: *ISG20*-KO spheroids. C: Counts per CAF cell of the of the top 5 most abundant UGIs in selected spheroid. Left: *AAVS1*-KO spheroids; right: *ISG20*-KO spheroids.

**Figure S3:**
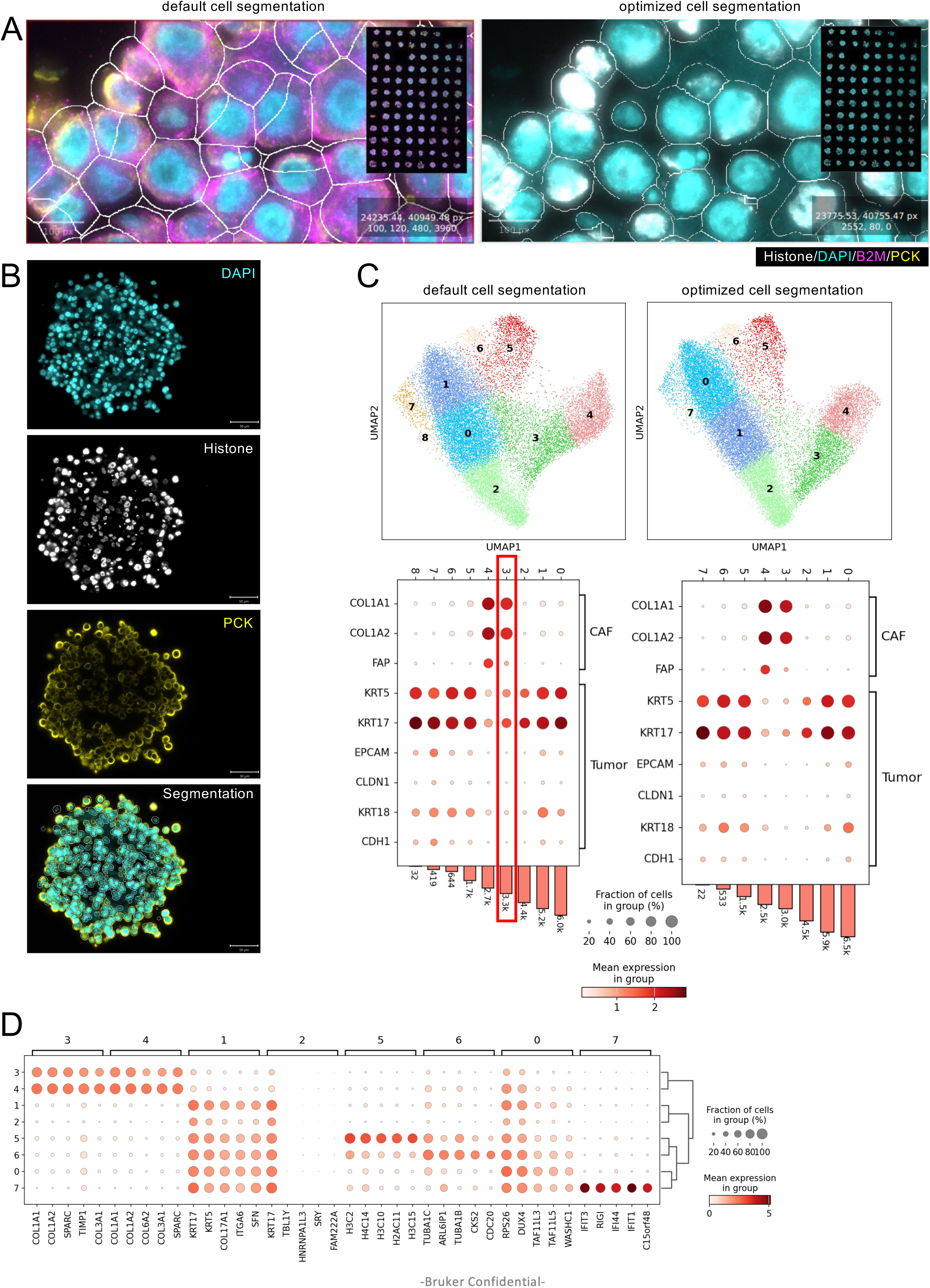
Optimized cell segmentation reduced doublets and improved cell type calling. A: Example of cell segmentation on the same region with default segmentation (left) and optimized segmentation (right). B: Example of optimized cell segmentation on a spheroid. Channels of DAPI, histone and PanCK protein staining were used for segmentation. Scale bar: 50 µm. C: Leiden clustering using default CosMx cell segmentation (left) and optimized cell segmentation (right), and expression level of cell type markers. The red box indicates cluster 3 in the default segmentation clustering, which shows high expression level of both fibroblast and tumor cell markers and thus indicating poorly segmented cells. Such mixed cell population is not seen with optimized cell segmentation. D: Expression level of marker genes in each cell cluster of CAF-tumor spheroid with optimized segmentation.

**Figure S4:**
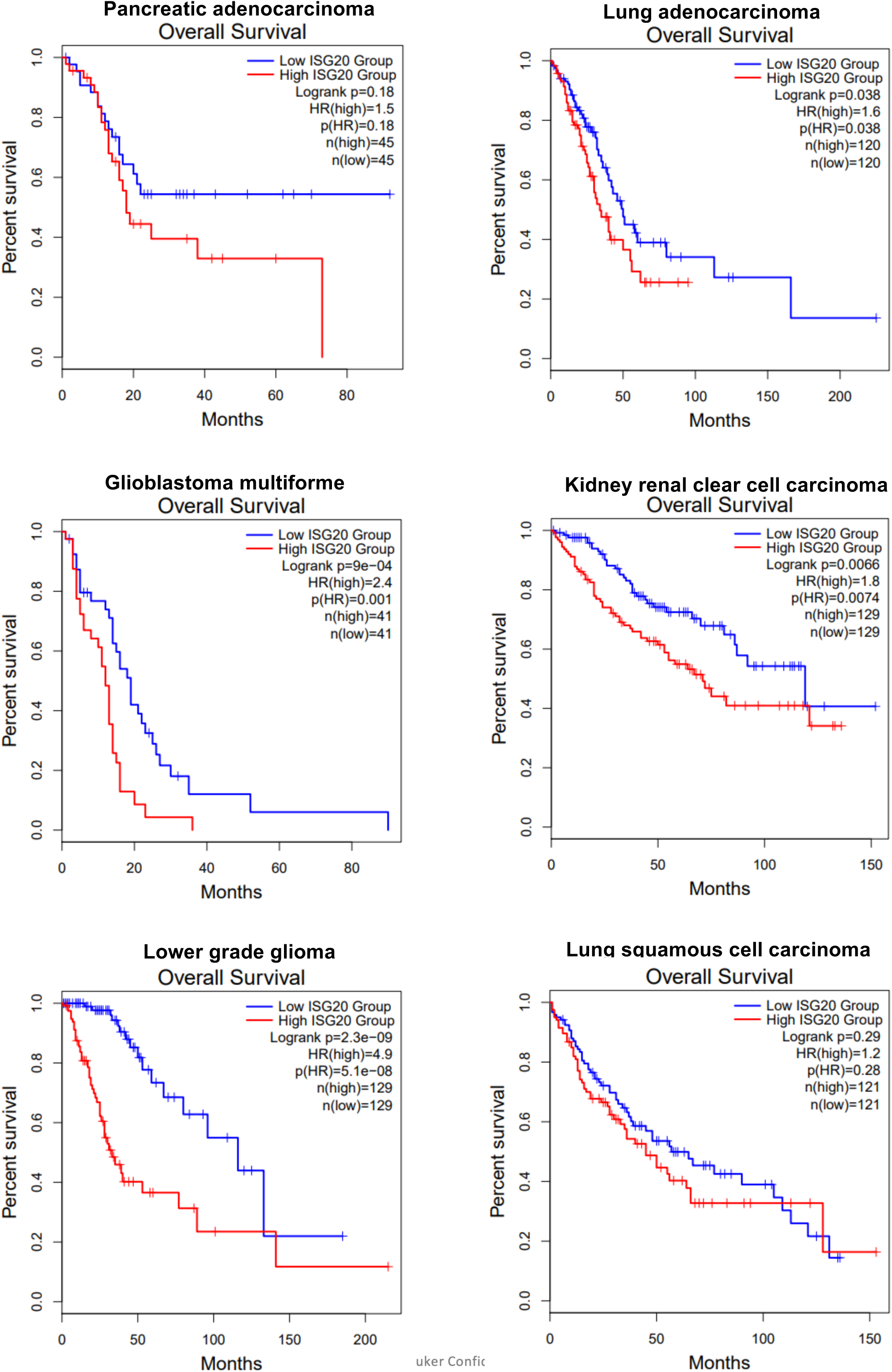
Patient survival in ISG20-high vs ISG20-low tumor types. Kaplan-Meier plots of patient survival in different tumor types that are ISG20-high and ISG20-low (Data source: TCGA/GTEx).

**Figure S5:**
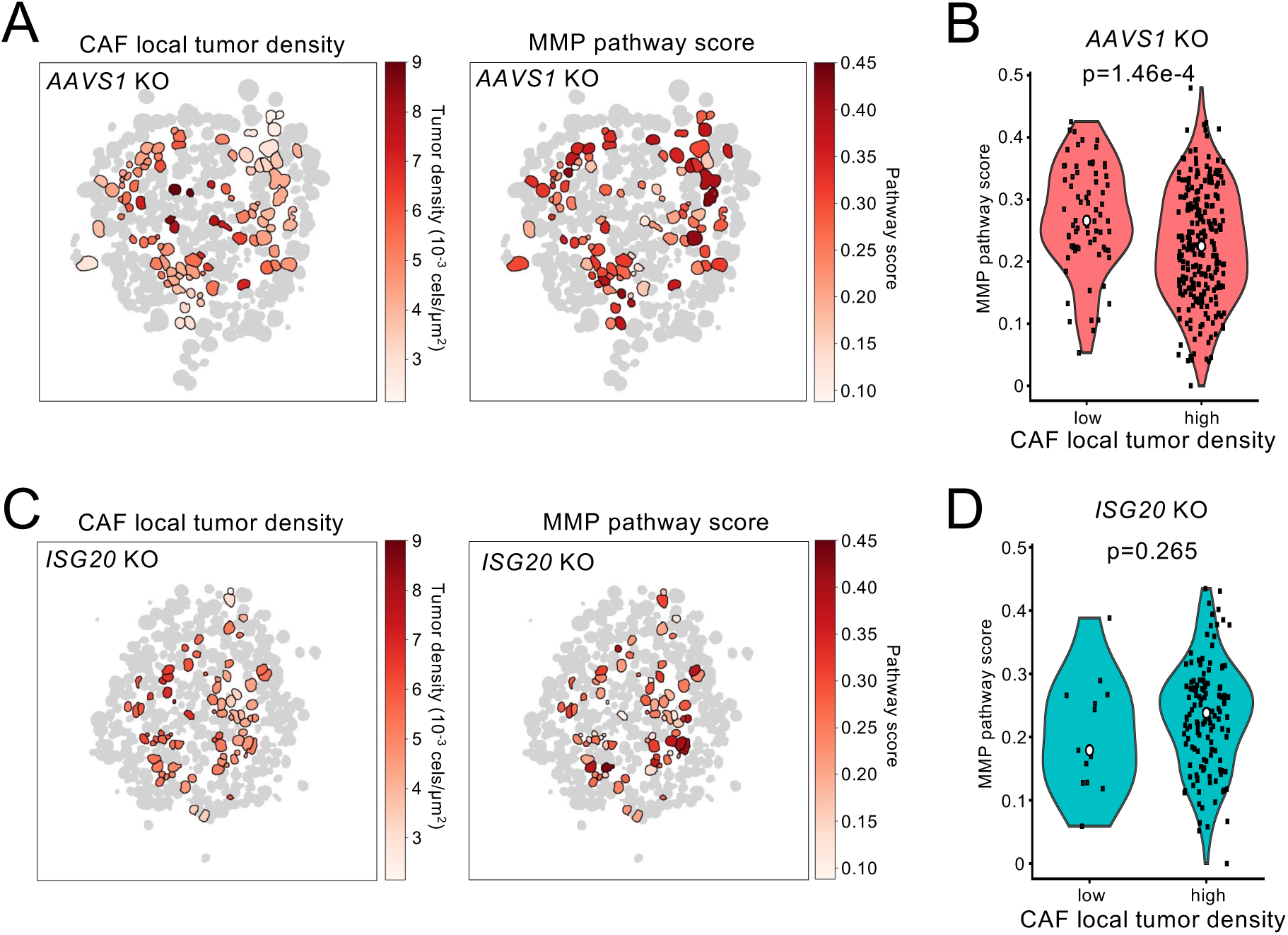
*ISG20* KO led to altered spatial activity of MMP pathway in CAFs. A: Heatmaps of local tumor density at CAF (left) and CAF MMP pathway activity (right) in *AAVS1*-KO spheroids. Tumor cells are shown in gray. B: Violin plot of CAF MMP pathway activity in *AAVS1*-KO spheroids, comparing CAFs with low tumor density (<0.003 tumor cells/µm^2^) versus high tumor density. P value was calculated by Wilcoxon two-sided rank-sum test. C: Heatmaps of local tumor density at CAF (left) and CAF MMP pathway activity (right) in *ISG20*-KO spheroids. Tumor cells are shown in gray. D: Violin plot of CAF MMP pathway activity in *ISG20*-KO spheroids, comparing CAFs with low tumor density versus high tumor density. P value was calculated by Wilcoxon two-sided rank-sum test.

**Figure S6:**
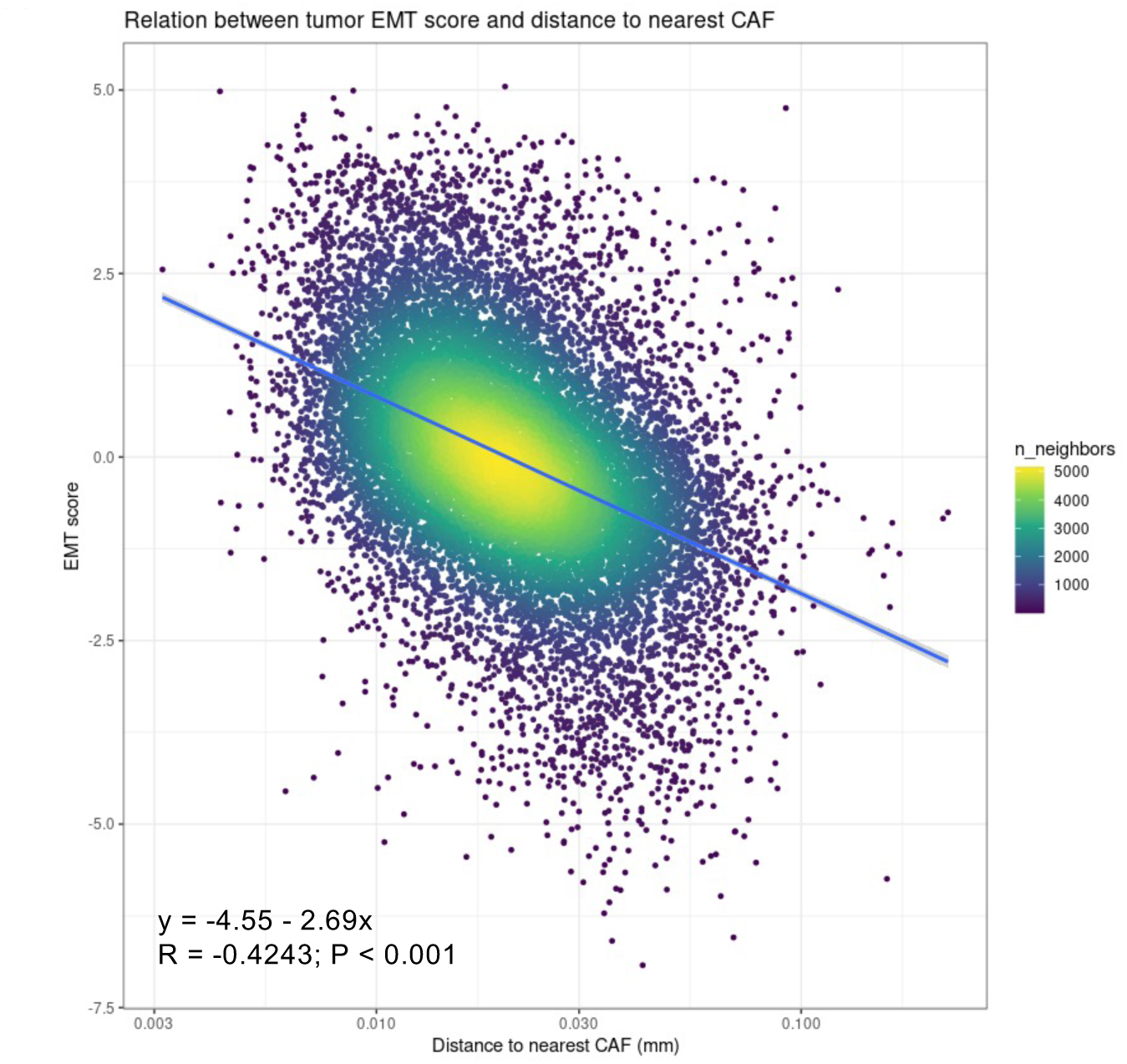
Tumor cell EMT score negatively correlates with their distance to the closest CAF cell. Correlation between tumor cell EMT score and their distance to the closest CAF cell.

**Figure S7:**
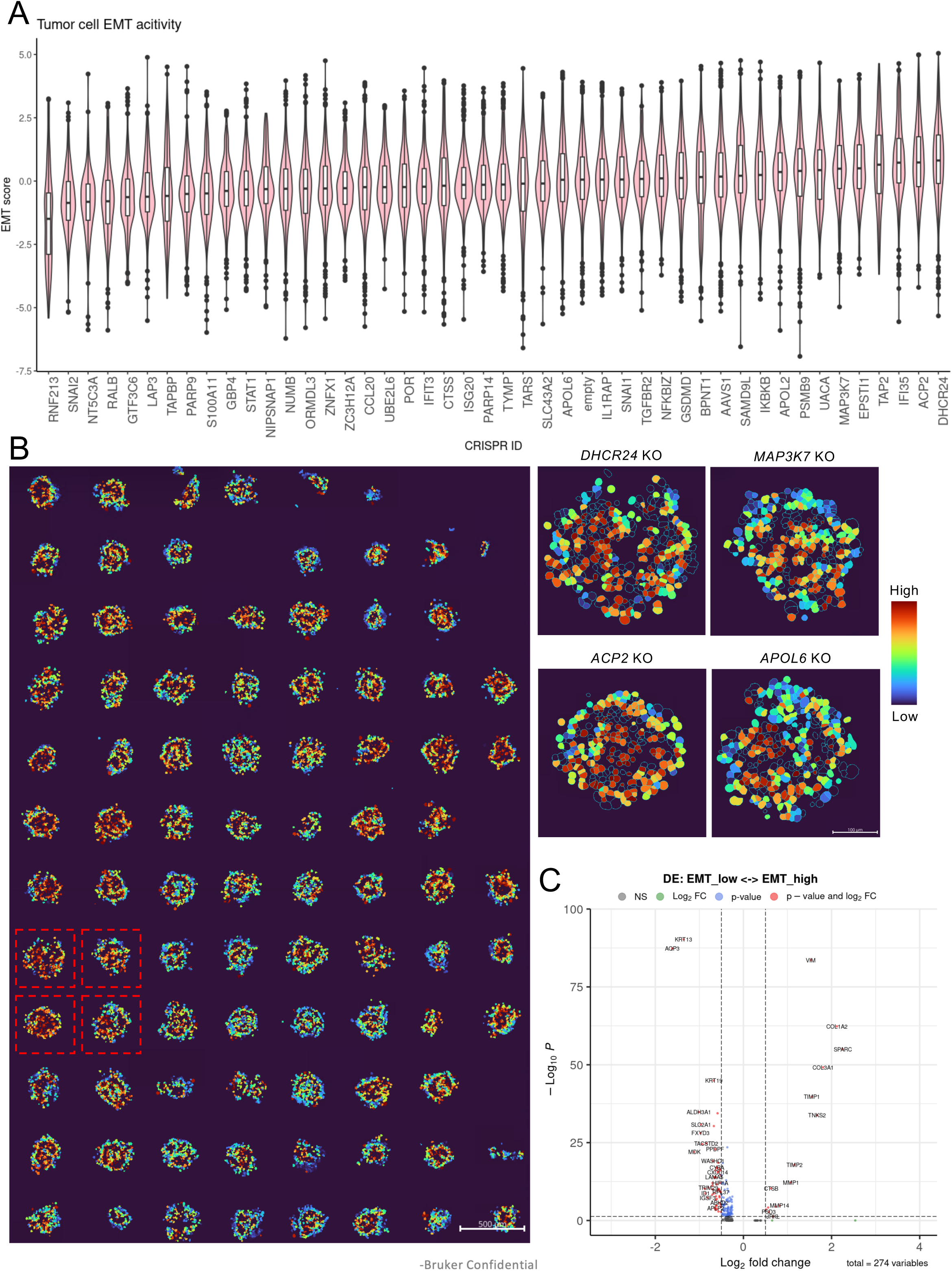
Multiomic SPACE enables unbiased analysis of tumor EMT score. A: EMT score of tumor cells co-cultured with different CAF KO cells, ranked from low to high from left to right. Tumor EMT score is calculated with normalized protein expression values: *EMT_score = f(Vimentin+SMA+Fibronectin) – f(EpCAM)* (also see Methods). B: Spatially resolved heatmap of EMT score in each individual tumor cells. Scale bar: 500 µm (left) and 100 µm (right). C: Differential gene analysis between tumor cells with high EMT scores (>Q3) and low EMT scores (<Q1).

**Table S1:** Comparison between SPACE and other single cell/spatial perturbation technologies.

|  | SPACE | Perturb-seq | Perturb-FISH | Perturb-Multi | RAEFISH |
| --- | --- | --- | --- | --- | --- |
| RNA coverage | Whole transcriptome | Whole transcriptome | ~ 130 genes | ~ 200 genes | ~ 300 genes |
| Same slide/cell protein coverage | 68 (Up to 76 targets) | Not demonstrated | Not demonstrated | 14 | Not demonstrated |
| Spatial resolution | Subcellular | No spatial info | Subcellular | Subcellular | Subcellular |
| Model complexity Demonstrated | 2D & 3D, FFPE compatible | Dissociated cells | Cultured cells | Fresh frozen tissues | Cultured cells |

Data as of Jan 16, 2026

**Table S2:** gRNA and UGI sequences for CosMx CRISPR identity detection in lentivirus-infected cell pool.

**Table S3:** gRNAs and UGI sequences for SPACE in CAF-tumor spheroids and their corresponding PCR primer sequences for CRISPR editing efficiency confirmation via NGS.

**Table S4:** CosMx probe design of CRISPR identity detection in lentivirus-infected cell pool.

**Table S5:** SPACE probe design of CRISPR identity detection in spheroids.

## Acknowledgements

We thank Heather Zhou (Merck & Co., Inc., Rahway, NJ, USA), Meagan Sullender (Merck & Co., Inc., Rahway, NJ, USA), the Broad Functional Genomics Consortium and UMass Medical School Gene Therapy Center for their help in designing the lentivirus vectors and generating lentiviruses. We thank Erin Piazza (Bruker) for help with RNA ISH probe design.

## Author contributions

conceived and designed the study: V.P.

performed experiment: M.H., S.M., K.C., M.A. with the help from S.I. CosMx CRISPR guide/barcode Probe designed by T.R. UGIs designed by D.B. and J.L.

analyzed the data: Y.C., Q.H., M.P. with the help from B.M., N.J., S.V.

supervised the study: V.P., S.H., J.B., A.T., M.R., M.S., F.P., I.K.

wrote the manuscript: M.H., Y.C., V.P. (with input from all authors).

## Declaration of generative AI and AI-assisted technologies

During the preparation of this work, the authors used ChatGPT (OpenAI) and Grok (xAI) in order to correct grammar mistakes and improve fluency of the manuscript. After using the tools, the authors reviewed and edited the content as needed and take full responsibility for the content.

